# Simulating big mechanically-active culture systems (BigMACS) using paired biomechanics-histology FEA modelling to derive mechanobiology design relationships

**DOI:** 10.1101/2024.12.08.627430

**Authors:** Sabrina Schoenborn, Mingyang Yuan, Cody A. Fell, Chuanhai Liu, David F. Fletcher, Selene Priola, Hon Fai Chan, Mia Woodruff, Zhiyong Li, Yi-Chin Toh, Mark C. Allenby

## Abstract

Big mechanically-active culture systems (BigMACS) are promising to stimulate, control, and pattern cell and tissue behaviours with less soluble factor requirements, however, it remains challenging to predict if and how distributed mechanical forces impact single-cell behaviours to pattern tissue. In this study, we introduce a centimetre, tissue-scale, finite element analysis (FEA) framework able to correlate sub-cellular quantitative histology with centimetre-scale biomechanics. Our framework is relevant to diverse bigMACS; media perfusion, tensile-stress, magnetic, and pneumatic tissue culture platforms. We apply our framework to understand how the design and operation of a multi-axial soft robotic bioreactor can spatially control mesenchymal stem cell (MSC) proliferation, orientation, differentiation to smooth muscle, and extracellular vascular matrix deposition. We find MSC proliferation and matrix deposition correlate positively with mechanical stimulation but cannot be locally patterned by soft robot mechanical stimulation within a centimetre scale tissue. In contrast, local stress distribution was able to locally pattern MSC orientation and differentiation to smooth muscle phenotypes, where MSCs aligned perpendicular to principal stress direction and expressed increased α-SMA with increasing 3D Von Mises Stresses from 0 to 15 kPa. Altogether, our new biomechanical-histological simulation framework is a promising technique to derive the future mechanical design equations to control cell behaviours and engineer patterned tissue generation.

## 1. Introduction

The engineering of lab-grown cell and tissue products depends, in part, on the mechanical environment that cells are exposed to [1]. There have been decades of research into the negative effects of mechanical stress; how the stiffness of different cell culture substrates alters the potency or differentiation of cells [2, 3], or how media perfusion imparts shear stresses that can impair cell health during culture [4, 5]. More recently, big mechanically-active culture systems (bigMACS) have been engineered to improve lab-grown cell and tissue product biomanufacturing [6, 7]. BigMACS include perfusion bioreactors to up-scale manufacturing [8], tensile stress bioreactors to improve myogenic differentiation [9], and soft robotic bioreactors whose on-demand flexion can enable non-invasive cell harvest [10]. BigMACS platforms are promising to enhance cell and tissue biomanufacturing but, to date, remain limited by inconsistent outcomes and an incomplete understanding of how their mechanical forces influence cell behaviours.

The impact of MACS mechanics on cell behaviour is typically evaluated by finite element analysis (FEA), with computational modelling correlated to tissue histology [11–13], as summarised in **Table 1**. A major shortcoming of this approach is that FEA approaches are often based on continuum mechanics theory focussed on tissue-scale analysis. These existing FEA approaches do not capture how local mechanical forces influence single-cell behaviours [14]. Cellular mechano-transduction depends on each cell’s local stresses (single-cell mechano-transduction), as well as the mechanics and actions that neighbouring cells experience (in paracrine mechano-transduction) [15]. Furthermore, MACS platforms often contain mechanical or structural artefacts at the microscale (such as microroughness caused during fabrication processes) [16], too miniscule to capture at the millimetre-to-centimetre tissue scale, which can greatly affect the local mechanics that a cell senses [17]. If we want to know how much mechanical stress to apply, to a specific region of our culture, at a specific time in culture, to control cell behaviour and to pattern lab-grown tissue, we must create new single-cell-scale mechanical models which capture the single cell micro-scale, as well as tissue mesoscale relevant to mechano-transduction.

**Table 1:** Effect of cyclic tensile strain on human mesenchymal stem cells (hMSCs) . ALP – alkaline phosphatase; αSMA - α-smooth muscle actin; or BMP-2 - bone morphogenetic protein 2; Cbfa1 - core binding factor α1; FGF-2 - fibroblast growth factor 2; IL1RN - interleukin-1 receptor antagonist; Runx2 - Runt-related transcription factor 2; PDMS – Polydimethylsiloxane; SM22α - smooth muscle 22α; VEGF-A – vascular endothelial growth factor A.

| Author, Year, [Reference] | Culture Device | Actuation Regime | Cell Behaviour |
| --- | --- | --- | --- |
| Chen et al., 2008, [13] | Collagen-I coated scaffolds | 3% cyclic tensile strain, 8h. | Upregulation of osteoblastic markers, increased levels of ALP and Cbfa1. Cell alignment perpendicular to axis of strain. |
|  |  | 10% cyclic tensile strain, 48h. | Increase in type collagen I, collagen III, and tenascin-C, indicating tenogenic differentiation. Cell alignment perpendicular to axis of strain. |
| Park et al., 2004, [18] | Elastin-coated membranes | 10% cyclic tensile strain, 24h, 1 Hz. | Transient increase in collagen I, α-actin, SM-22α, and β-actin. Cell alignment perpendicular to axis of strain. |
| | | 10% cyclic tensile strain, 3 days, 1 Hz. | SM-22α decreased by 25% after alignment. Cell alignment perpendicular to axis of strain. |
| | Collagen-coated membranes | 10% cyclic tensile strain, 24h, 1 Hz. | Transient increase in collagen I, α-actin, SM-22α, and β-actin. Cell alignment perpendicular to axis of strain. |
| | | 10% cyclic tensile strain, 24h, 1 Hz. | 10% cyclic tensile strain, 3 days, 1 Hz SM-22α decreased by 50% after alignment. Cell alignment perpendicular to axis of strain. |
| **Khani et al., 2015, [19]** | Collagen-coated PDMS. | 5% cyclic tensile strain, 24h, 1 Hz. | Strained cells without TGF-β1 significantly increased Young's Moduli and elevated levels of smooth muscle markers αSMA, h1-Calponin and SM 22A. Strained hMSCs demonstrated increased myogenesis. Cell alignment perpendicular to axis of strain. |
| **Koike et al., 2005, [20]** | Collagen-I coated membranes | 0.8% cyclic tensile strain, 48h, 1 Hz. | Cbfa1/Runx2 increased. ALP activity increased. |
|  |  | 5% cyclic tensile strain, 48h, 1 Hz. | Cell proliferation significantly increased. Cbfa1/Runx2 increased. Collagen I increased. Osteocalcin expression and ALP activity significantly decreased. |
|  |  | 10% cyclic tensile strain, 48h, 1 Hz. | Cell proliferation significantly increased. Collagen I increased. Osteocalcin expression and ALP activity significantly decreased. |
|  |  | 15% cyclic tensile strain, 48h, 1 Hz. | Cell proliferation significantly increased. Cbfa1/Runx2 decreased. Collagen I Increased. Osteocalcin expression and ALP activity significantly decreased. |
| **Jagodziniski et al., 2004, [21]** | Silicone scaffold. | 2% cyclic tensile strain, 6h/day for 3 days, 1 Hz. | Increased ALP secretion and collagen III upregulation. Enhanced hMSC osteogenic commitment. |
|  |  | 8% cyclic tensile strain, 6h/day for 3 days, 1 Hz. | Increased ALP secretion and collagen III upregulation. Collagen I and Cbfa1 upregulated. Enhanced hMSC osteogenic commitment. |
| **Sumanasinghe et al., 2006, [22]** | 3D collagen matrix. | 10% cyclic tensile strain, 4h/day for 7 and 14 days, 1 Hz. | Increase in BMP-2 expression for both durations of strain. |
|  |  | 12% cyclic tensile strain, 4h/day for 7 and 14 days, 1 Hz. | Increase in BMP-2 expression only in cells subjected to 14 days of strain. |
| **Charoenpanich et al., 2011, [23]** | Collagen I gel sheet | 10% cyclic tensile strain, 4h/day for 14 days, 1 Hz. | Osteogenic differentiation, upregulation of IL1RN, FGF-2, VEGF-A. |
| **Qiu et al., 2016, [24]** | Collagen fibres | 10% cyclic tensile strain, 12h/day for 14 days, 1 Hz. | Collagen I, collagen III, tenascin-C, and fibroblastic transcription factor scleraxis upregulated. Tenogenic differentiation. |
| **Yang et al., 2012, [25]** | PEG hydrogels | 10% cyclic tensile strain, 12h/day for 14 days, 1 Hz. | Upregulated collagen III and upregulated tenascin-C. No cell alignment observed. |

In 2022, our group published details of a soft robotic bioreactor for mesenchymal stem cell (MSC) culture [26]. We designed our soft robotic bioreactor to model the radial and angular multi-axial mechanical forces found within the human femoropopliteal artery while standing, walking, and crouching in normal and diseased cases. We found increasing mechanical forces with values similar to those in our body’s normal femoropopliteal artery led to increased MSC proliferation, alignment, and smooth muscle cell (SMC) differentiation in the absence of soluble differentiation factors. Although limited to bulk tissue analysis, local MSC behaviours appeared to be patterned by the quantity and angle of the mechanical forces we applied in our soft robotic bioreactor, as also observed by other studies using pneumatic and soft robotic MACS platforms with distributed mechanical forces [10, 27–30]. If we could define a predictive relationship between mechanical forces and patterned tissue generation, we could engineer more realistic and functional lab-grown cell and tissue products.

In this paper, we develop a biomechanics-histology framework for the simulation of big mechanically-active culture systems (BigMACS) as applied to our prior soft robotic femoropopliteal artery bioreactor. Our approach utilises a FEA mesh of an extremely high resolution (10^6-7^ nodes/cm^2^), 100 to 1,000-fold more dense than a confluent cell monolayer, necessary to investigate the effect of mechanical forces on single-cells on a 3D printed scaffold. This FEA mesh resolution is essential as microrough surfaces are often unavoidable in bigMACS and 3D printed platforms, and microrough surfaces play important roles in single-cell mechano-transduction [31]. Scaling our single-cell-scale simulations to tens of thousands of cells within centimetres of mechanically-stimulated tissue, we quantitatively understand whether and how local mechanical stress influence cell density, orientation, differentiation (MSCs to SMCs), and matrix deposition (collagen type-4). Our models indicate several of these mechanically-directed cell behaviours appear to occur or stop once a threshold stress magnitude is reached.

This paper proposes a single-cell biomechanics-histology predictive modelling framework which seeks to uncover fundamental design equations in mechanobiology at scales relevant to the nascent cell and tissue biomanufacturing industry. We believe these new approaches could have a broad impact across cell and tissue types and small to big mechanically active culture systems (BigMACS).

## 2. Methodology

### 2.1. Fabricating the BigMACS Soft Robotic Femoropopliteal Artery Bioreactor

Soft robotic bioreactors were engineered within a previous study [26]. Briefly, two-part soft robotic bioreactor moulds were printed from 3D Tuff resin (Monocure 3D, Sydney, Australia) using a digital light projection printer (DLP, Creality LD-002R, Shenzhen, China), washed, and further cured. Platsil OO-20 silicone was poured into the top and bottom moulds and placed onto a hotplate at 60°C for at least 2 hours until solidified. Two-part Platsil OO-20 silicone casts were then adhered to one another with a thin layer of Platsil OO-20 silicone and placed back onto the hotplate for an additional 2h.

A rectangular recess in the soft robotic BigMACS bioreactor was coated with 10 μg/mL fibronectin and then seeded at 3 × 10^4^ TeloASC52 MSCs/cm^2^ (American Type Culture Collection) and expanded for 3 days until confluent. Then, soft robotic BigMACS bioreactors were pressurised by a custom rack-and-pinion syringe pump to undergo static, angular, radial, or angular and radial stress cycles every 1- 2 s for 24 hours (roughly 50,000 cycles), as described in **Table 2**. Cultures were then fixed with 4% paraformaldehyde (PFA), stained with rabbit anti-human collagen type 4 (col-IV), goat anti-human α- smooth muscle actin (α-SMA), phalloidin to stain cytoskeletal actin, and 4′,6-diamidino-2- phenylindole (DAPI) to stain cell nuclei. Soft robotic BigMACS were then imaged by a Nikon A1R confocal microscope (Tokyo, Japan).

**Table 2:**
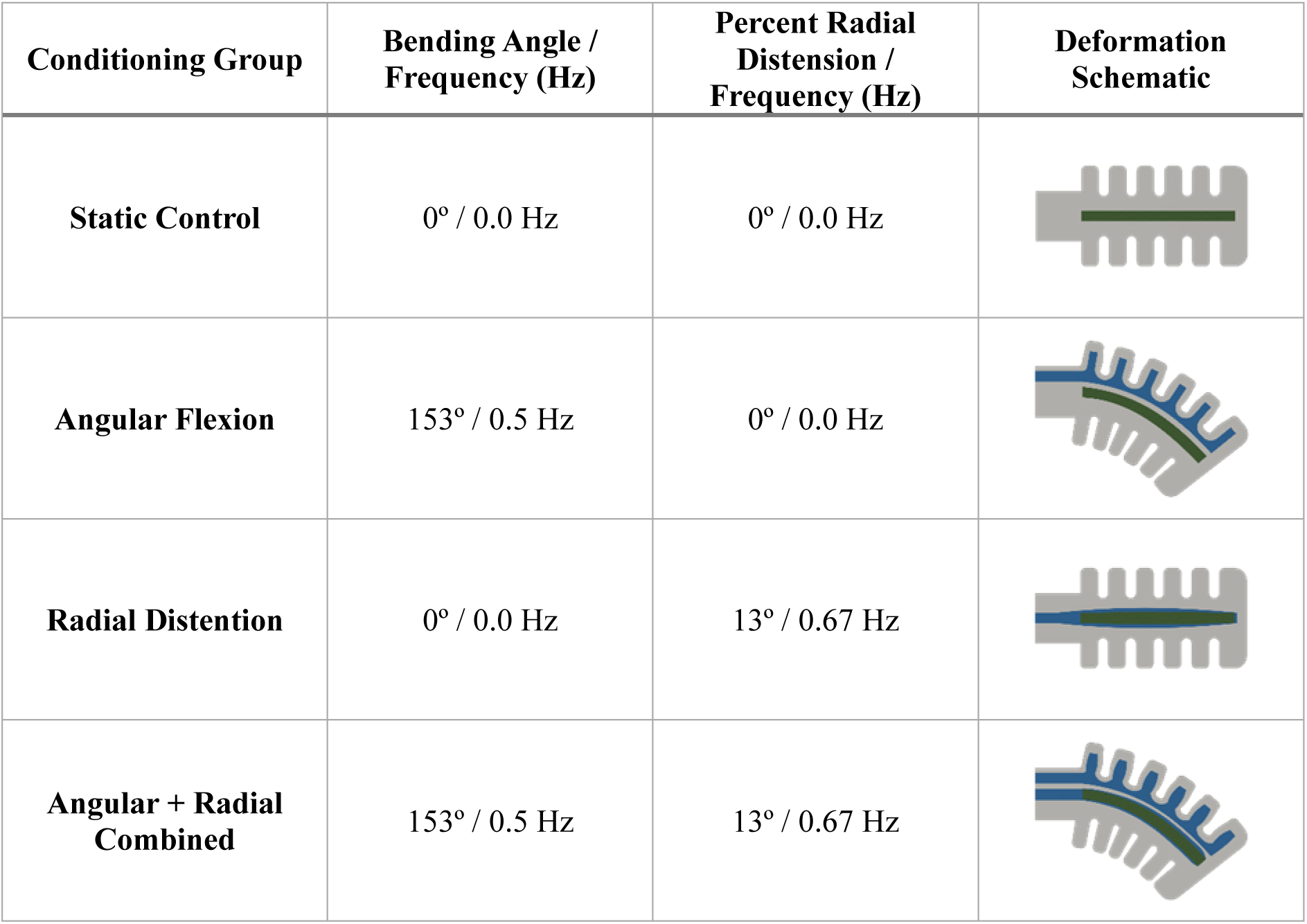
BigMACS mechanical stimulation conditions. Previously, four mechanical conditions were compared with three or four biological replicates each. Conditions included a static control, angular flexion, radial distension, and angular flexion and radial dissension combined. Bending angle and frequency replicated the angle and frequency of mechanical stimulation in the human femoropopliteal artery while walking. This figure is modified with permission from [26].

| Conditioning Group | Bending Angle / Frequency (Hz) | Percent Radial Distention / Frequency (Hz) | Deformation Schematic |
| --- | --- | --- | --- |
| Static Control | 0° / 0.0 Hz | 0° / 0.0 Hz |  |
| Angular Flexion | 153° / 0.5 Hz | 0° / 0.0 Hz |  |
| Radial Distention | 0° / 0.0 Hz | 13° / 0.67 Hz |  |
| Angular + Radial Combined | 153° / 0.5 Hz | 13° / 0.67 Hz |  |

### 2.2. BigMACS Finite-Element Analysis (FEA) ANSYS Simulation

Due to the large deformation of the soft-robot particularly during bending, a hyperelastic third-order Ogden model was chosen to describe the PlatSil® Gel-OO20 (Polytek) silicone material model [32, 33]:

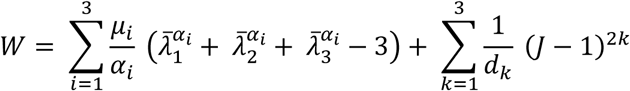

where 𝑊 is the strain energy potential, 𝜆̅_1_, 𝜆̅_2_, 𝜆̅_3_ are the deviatoric principal stretches, defined as 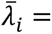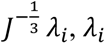 is the principal stretches of the left Cauchy-Green tensor, 𝐽 is the determinant of the elastic deformation gradient, and 𝜇_𝑖_, 𝛼_𝑖_ and 𝑑_𝑘_ are material constants. The initial shear modulus 𝜇 is defined:

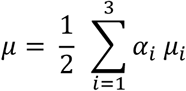

The Ogden model parameters were determined using the Ansys hyperelastic curve fitting function on uniaxial tensile data generated with the manufactured dog bone samples and biaxial and compressibility data from literature [34], with initial bulk modulus 𝐾 = 2⁄𝑑_1_, as summarised in **Table 3**.

**Table 3:** Third-order Ogden model parameters of PlatSil soft robot model. . Incompressibility parameters d_i_, material constants α_i_, material constants μ_i_.

| $\mu_i$ [MPa] | $\alpha_i$ [-] | $d_i$ [MPa<sup>-1</sup>] |
| --- | --- | --- |
| $\mu_1 = 1.4736 \times 10^{-2}$ | $\alpha_1 = 3.41576$ | $d_1 = 3.2587$ |
| $\mu_2 = 6.6703 \times 10^{-5}$ | $\alpha_2 = 7.0679$ | $d_2 = 0$ |
| $\mu_3 = 4.5381 \times 10^{-4}$ | $\alpha_3 = -3.3659$ | $d_3 = 0$ |

All 3 actuation groups of the original study were simulated by FEA: radial distension (RD) conditioning with 13% stretch cyclically at 0.67 Hz, angular flexion (AF) conditioning with a 153° angle of actuation cyclically at 0.5 Hz and a combined angular-radial (AR) conditioning. The cyclic actuation pressure provided by a syringe pump in the original study was modelled as a pressure boundary condition acting on the internal pneumatic channels and the resulting deformation of the soft robot model was analysed to ensure it matched the deformation of the experimental devices. Rigid body motion was prevented by applying a fixed support to the base area of the model, which was constrained by a rigid casing connecting it to the syringe pump in the experimental setup. The effect of gravity was included. A timestep was considered as converged when the force convergence value and the displacement convergence value were both below a residual target of 10^-4^. The model was run on the University of Queensland’s Bunya supercomputer [35] as a transient analysis using the sparse direct solver with a time-step size of 10 ms ensuring solver convergence and satisfying the timestep sensitivity analysis.

The maximum element size for the meshing of the cell culture area was 50 µm. While mesh convergence was achieved for larger element sizes, this value was chosen to facilitate a more detailed comparison between the histology imaging data and the numerical data since the average cell size of the MSCs utilised in this study was 50 µm. However, during imaging, the presence of microgrooves resulting from stereolithography (SLA) 3D printing layer lines were noted at an average periodicity of 105 µm tip-to-tip (**Figure 1A**). In the cell culture study, a small effect on cellular morphology and alignment was observed, which was not significant compared with the effect of the actuation. To facilitate a comparison of the local stress-strain profiles of both the ideal CAD geometry and the manufactured micro-grooved geometry with the histology imaging data, the original CAD model was modified to include these layer lines across the cell culture area of the model (**Figure 1A)**.

**Figure 1:**
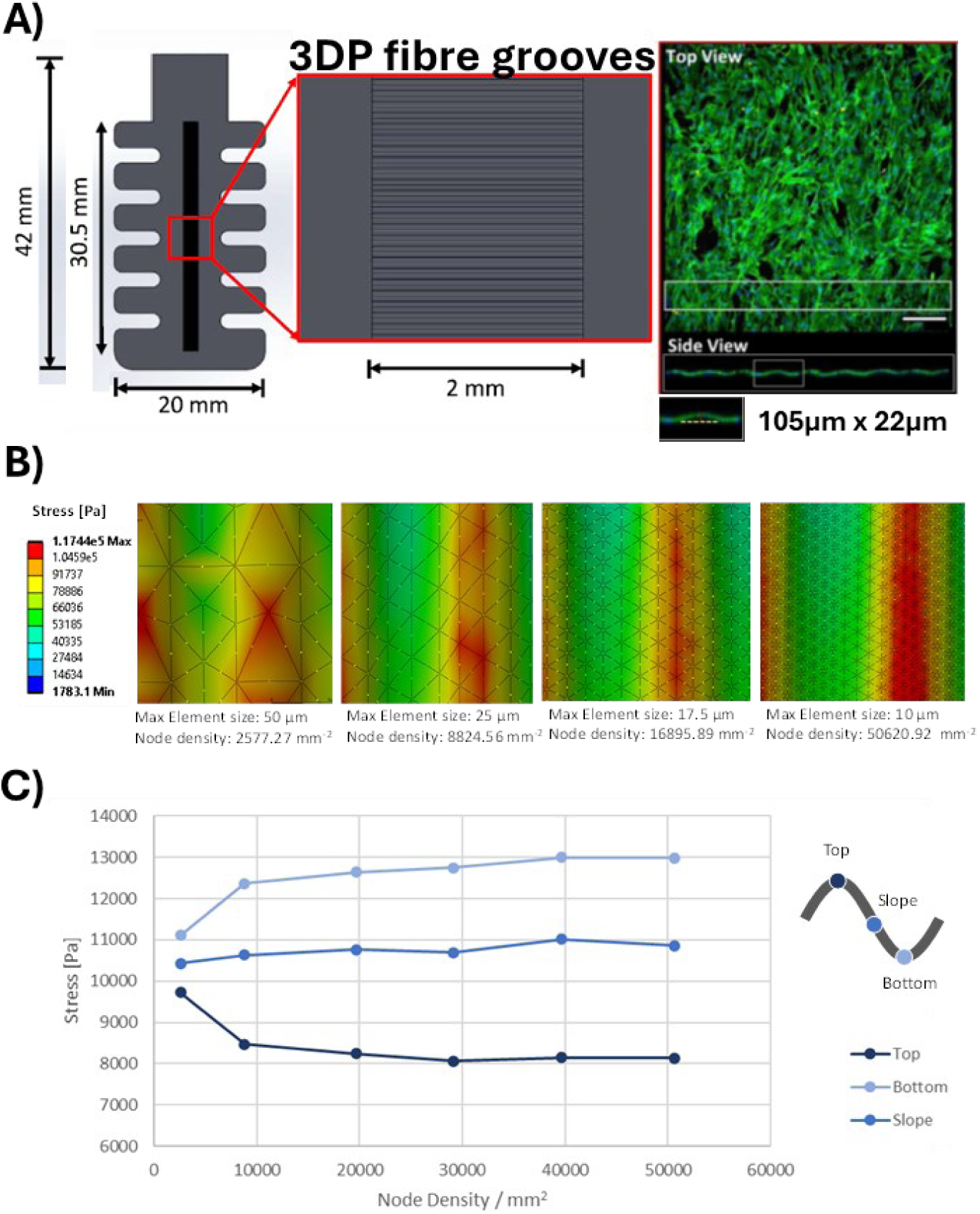
BigMACS geometry and FEA mesh sensitivity analysis. (A) Illustration of the bigMACS platform, showing the presence of microgrooves in the cell culture chamber, with confocal imagery of the 22 µm tall groove height. (B) Mechanical stress results of micro-grooved bigMACS under radial distension with various FEA mesh resolutions, showing inconsistent stress profiles below 10,000 mesh nodes per mm^2^. (D) Sensitivity of FEA mesh node density on Von Mises stress at top, bottom, and mid-point slope areas. Subfigure A is a modified version of supplemental data from [26].

This Ansys FEA simulation calculated all normal stress tensors (𝜎_𝑥_, 𝜎_𝑦_, 𝜎_𝑧_) and shear stress tensors (𝜏_𝑥𝑦_, 𝜏_𝑥𝑧_, 𝜏_𝑦𝑧_) for millions of simulated nodes. To estimate stress magnitude, we calculated 3D Von Mises Stress [36] from our FEA simulation’s node output as:

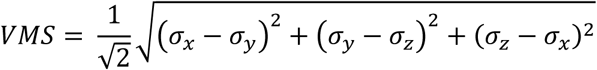

To estimate principal stress direction, we calculated 2D Von Mises Stress by setting the final dimension 𝜎_𝑖_ = 0. We also calculated angles of deviance from a principal stress tensor (Mohr’s Circle) in two dimensions (𝑖, 𝑗) [37, 38]:

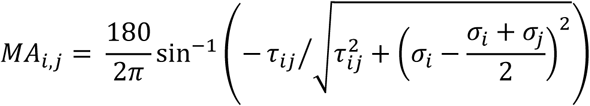

In the main figures of this paper, we focussed on 3D Von Mises Stress and angle on XY Mohr’s circle, but we have included all 2D Von Mises Stress and angle on XZ, YZ Mohr’s circle plots in the supplemental figures.

### 2.3. BigMACS Quantitative Histology Image Analysis and Statistics

The centre region of the soft robotic bigMACS culture recess was imaged by confocal microscopy by stitching together eight 3D image tiles (4 × 2 tilescan; roughly 20 mm^2^ of cell culture space) using NIS-Elements (Nikon). Col-IV, α-SMA, actin, and DAPI stains were detected with ImageJ’s particle analysis plugin, and actin cytoskeletal orientation was measured with ImageJ’s OrientationJ plugin [39]. These measurements were exported as comma-separated value spreadsheets with each row listing the location, intensity, and orientation of each Col-IV, α-SMA, actin, and DAPI stain, with 5,000 to 100,000 rows per stain per soft robotic bioreactor. These spreadsheets were imported into python to correlate each stain’s local density (stain/cm^2^) and orientation with the local principal stress magnitude and direction, specifically 3D Von Mises Stress and XY Mohr’s Angle, through a third order polynomial regression (Python’s polyfit) as presented alongside 95% confidence intervals in **Figures 4 and 5**. Measurement replicates calculated for the polynomial regression confidence intervals were considered as each individual stain in the image (with 5,000 to 100,000 stains detected per image); it is important to note that only one biological and one image replicate were compared per soft robotic bioreactor condition.

Local stain density (stain/cm^2^) was calculated for each stain by counting the number of stains within a 3D radius (e.g., 50 µm), projected to 2D by averaging the neighbour count for 𝑥 and 𝑦 pairs, and dividing by its circular area (7850 µm^2^). However, this approach creates edge effects when measuring local density for stains on the edge of the image, where part of the circle’s area is outside the imaging space, an image analysis challenge previously recognised [40]. To overcome this edge effect, we mirrored the distribution of cells on the opposite side of each of four 2D imaging edges (left, right, top, bottom). As the imaging edges were sometimes irregular (due to artefacts) we preformed this mirroring exercise for many discrete steps along each edge, outlined in **Supplemental Figure S1**.

## 3. Results and Discussion

### 3.1. Simulating BigMACS Requires Micro-Resolved FEA due to 3D Printing Micro-Artefacts

A mesh sensitivity analysis on the grooved soft robotic model indicates that the computational cost of resolving microgrooves in the model is considerable compared with resolving to a cellular level. Since the microgrooves generated via SLA printing (105 µm) are approximately twice the size of an average MSC (50 µm), the microgrooves cannot be sufficiently resolved using a mesh of this size and local stress gradients between the top and bottom of grooves are not resolved (**Figure 1B, C**). An element size of 25 µm (25% the size of the groove) allows for the resolution of a distinct stress gradient along the groves but it requires an element size of approximately 10 µm or 10% of the size of the microgroove to reach a mesh convergence target of below 2% resulting in an increase in node number of over 1960% for the micro-grooved cell culture region compared with a maximum element size of 50 µm.

The results obtained with a maximum element size of 10 µm demonstrates the significant effect of microgrooves on local stress values, with stresses at the top and bottom of grooves fluctuating 20-30% around the respective average stress on a flat surface. Where microgrooves need to be numerically resolved, the required node density in our study increased by a factor of 20 from a mesh-independent smooth model to a mesh-independent micro-grooved model, indicating that the elimination of microgrooves for in-vitro/in-silico coupled models is highly beneficial with regards to computational cost. Smooth bigMACS simulations required 100-fold fewer nodes to achieve mesh convergence (10^4^ instead of 10^6^ nodes/cm^2^), in-line with existing approaches. A comparison of micro-grooved versus smoothed soft robotic bioreactor Von Mises stress normal and shear stresses and Mohr’s Stress Angles are in **Supplemental Figures S2-S7**, while a comparison of micro-grooved and smoothed histological images can be found in our prior publication’s supplemental data [26]. The effect of micro-grooves were neglected as their effect appeared to be less significant than the effect of BigMACS actuation.

### 3.2. BigMACS Exhibit Highly Heterogenous Mechanical Stress Gradients

The stress simulations illustrated the highly heterogeneous mechanical stresses applied to the soft robotic bigMACS cell culture area (**Figure 2**). BigMACS undergoing radial distension exhibited a bulge in their cell culture area, turning the concave recess into a convex bulge, and applying maximum and average 3D Von Mises Stress of 14 kPa and 10 kPa, concentrated at the centre of the cell culture recess. Radial distension bioreactors exhibited symmetric XY Mohr’s Angle to the left and right of the culture recess, with XY Mohr’s Angles from −30 to +30 degrees.

**Figure 2:**
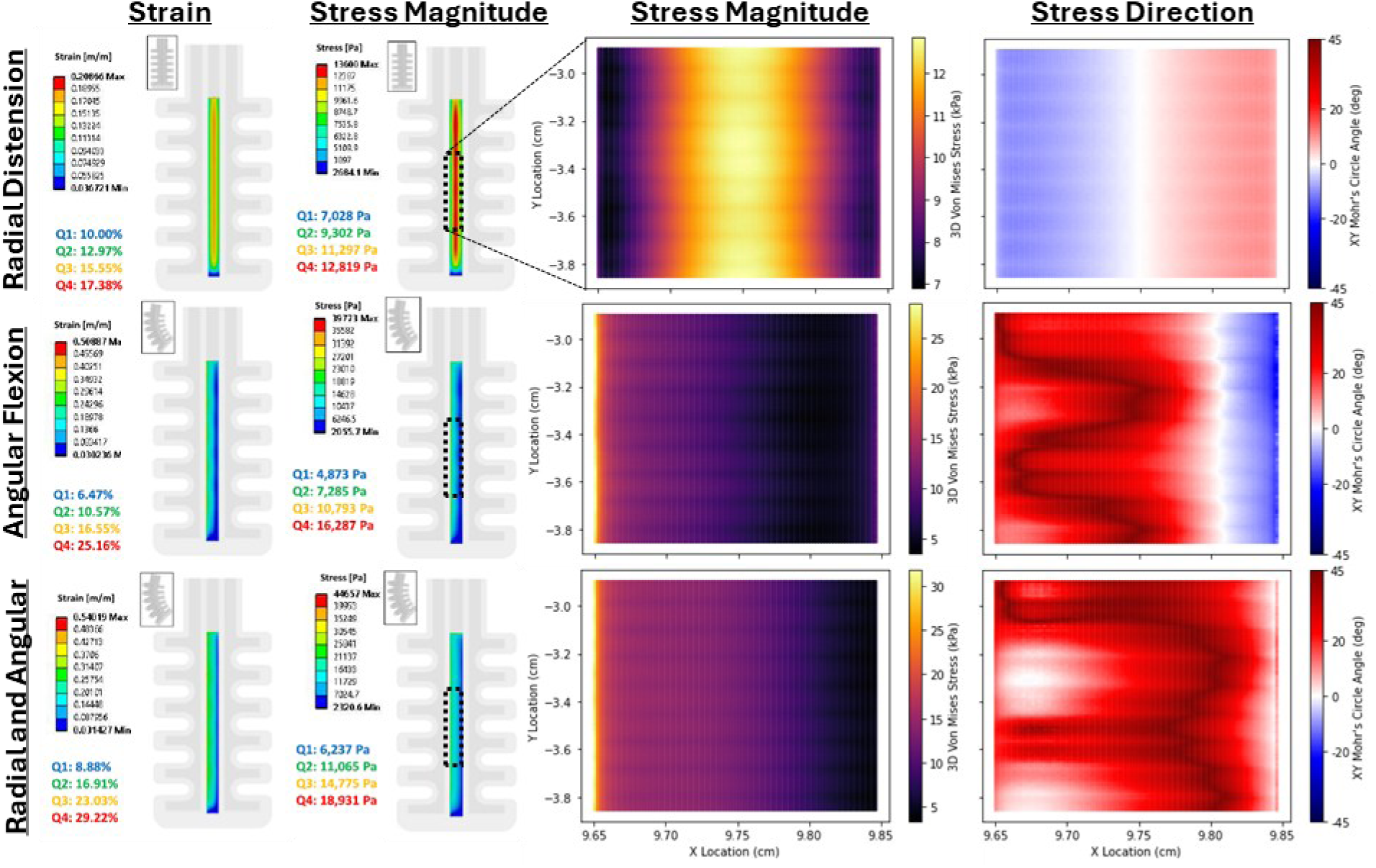
Biomechanics simulations of bigMACS. Full-scale strain and stress-magnitude simulations of bigMACS under radial distension, angular flexion, and radial and angular distension. (Right) Stress magnitude (3D Von Mises Stress) and stress direction (XY Mohr’s Angle) of a cell culture region imaged by confocal microscopy.

The bigMACS undergoing angular flexion exhibited a much higher maximum but lower average 3D Von Mises stress than radial distension (40 kPa, 8.5 kPa), concentrated on the outer arc of the bending culture recess (left side), which dissipated quickly when moving across the culture recess to the inner arc (right side). The angular flexion bioreactor’s XY Mohr’s Angle was greatly affected by the presence of microgrooves every 105 µm but also by the pneumatic chambers spaced every 500 µm on the sides of the soft robot. During angular flexion, the pneumatic chambers along the inner arc resist compression and create a stress oriented in the opposite direction on the right-hand side of the culture recess.

When combining radial distension with angular flexion, the maximum and average 3D Von Mises Stress are somewhat higher than angular flexion and radial distension, respectively (50 kPa, 13 kPa), and the soft robot’s compressed pneumatic chambers offered less resistance against angular flexion, exhibited by a consistently positive XY Mohr’s Angle. These simulations illustrate the complex microscale stress gradients impacting 3D printed and soft robotic bigMACS, due to microrough artefacts and pneumatic chambers ubiquitous to their production and designs, able to create significant impact at the cell scale.

### 3.3. BigMACS Conditioning Alters Cell Behaviours at Bulk and Local Scale

After four days of growth, or three days of growth and one day of mechanical conditioning, all four soft robot mechanical conditions exhibited confluent cell coverage across their culture recess (**Figure 3A**). Interestingly, all types of mechanical conditioning caused this confluent layer to become much more packed with cells, as quantitative histology analyses indicated that average nuclear density is five-fold higher for all mechanical conditioning regimes in comparison to the static control (**Figure 3B**). Nuclear density appeared to not be locally correlated with microscale stress magnitude, except for the highest 3D Von Mises Stresses from 15 to 20 kPa (**Figure 4**). The principal stress direction did not appear to correlate correlation to nuclear density. However, an angular flexion XY Mohr’s Stress Angle of −5°, where pneumatic channels along the soft robot’s inner arc began to push back against angular flexion (**Figure 3**), appear to correlate with a higher difference in nuclear density (**Figure 5**).

**Figure 3:**
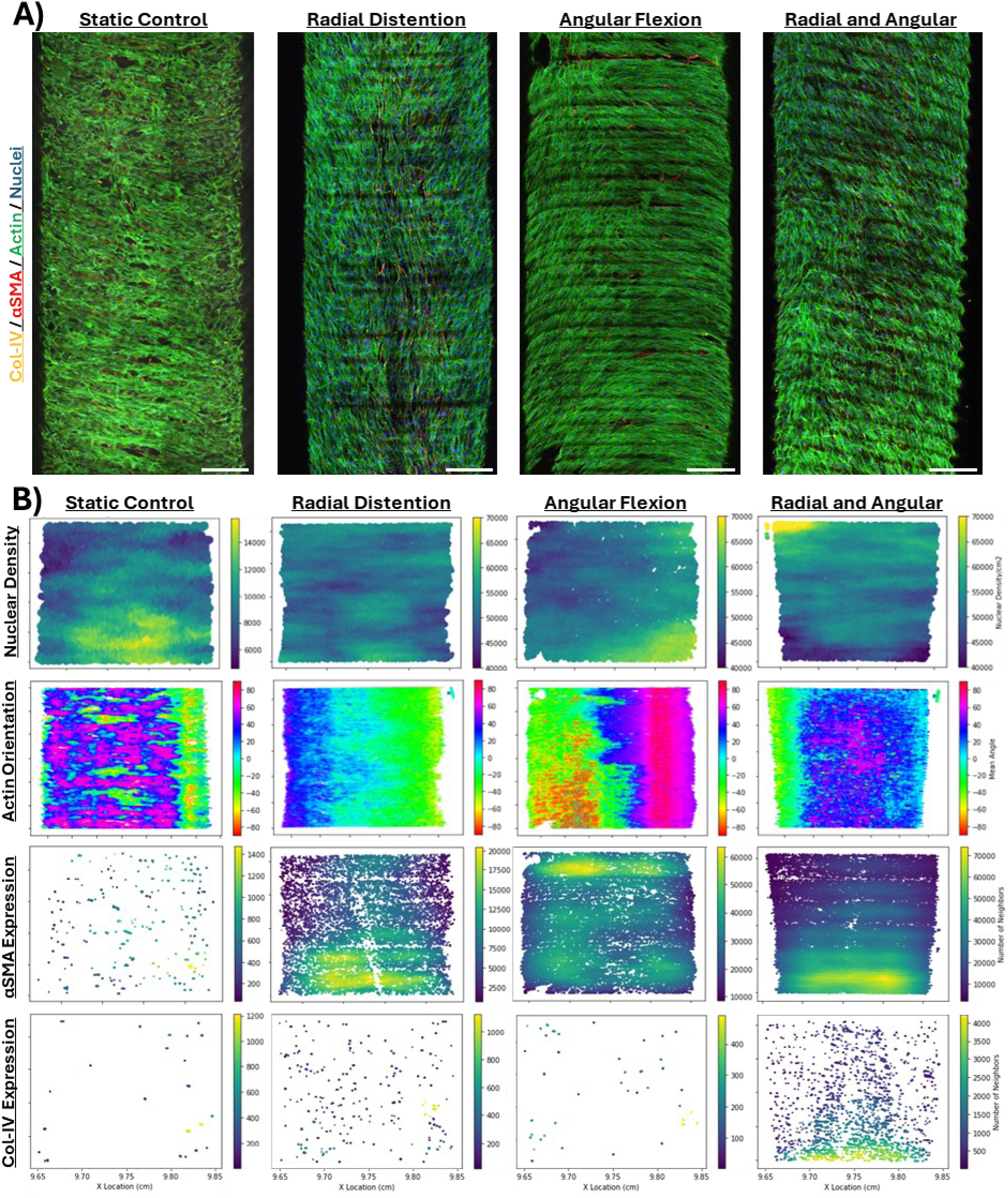
Quantitative histology of bigMACS conditioning. (A) MSCs were cultured in soft robotic bioreactors for four days in static (left), or for three days in static and one day in radial, angular, or radial and angular mechanical conditioning (right) and imaged by 4×2 tiled 3D confocal microscopy. Scale bars are 0.05 cm. (B) Computational histology analyses of the nuclear density, actin orientation, α-SMA stain density, and Col-IV stain density of the four bigMACS conditions (note that heatmap scale for the static control density analyses are several-fold lower than others).

**Figure 4:**
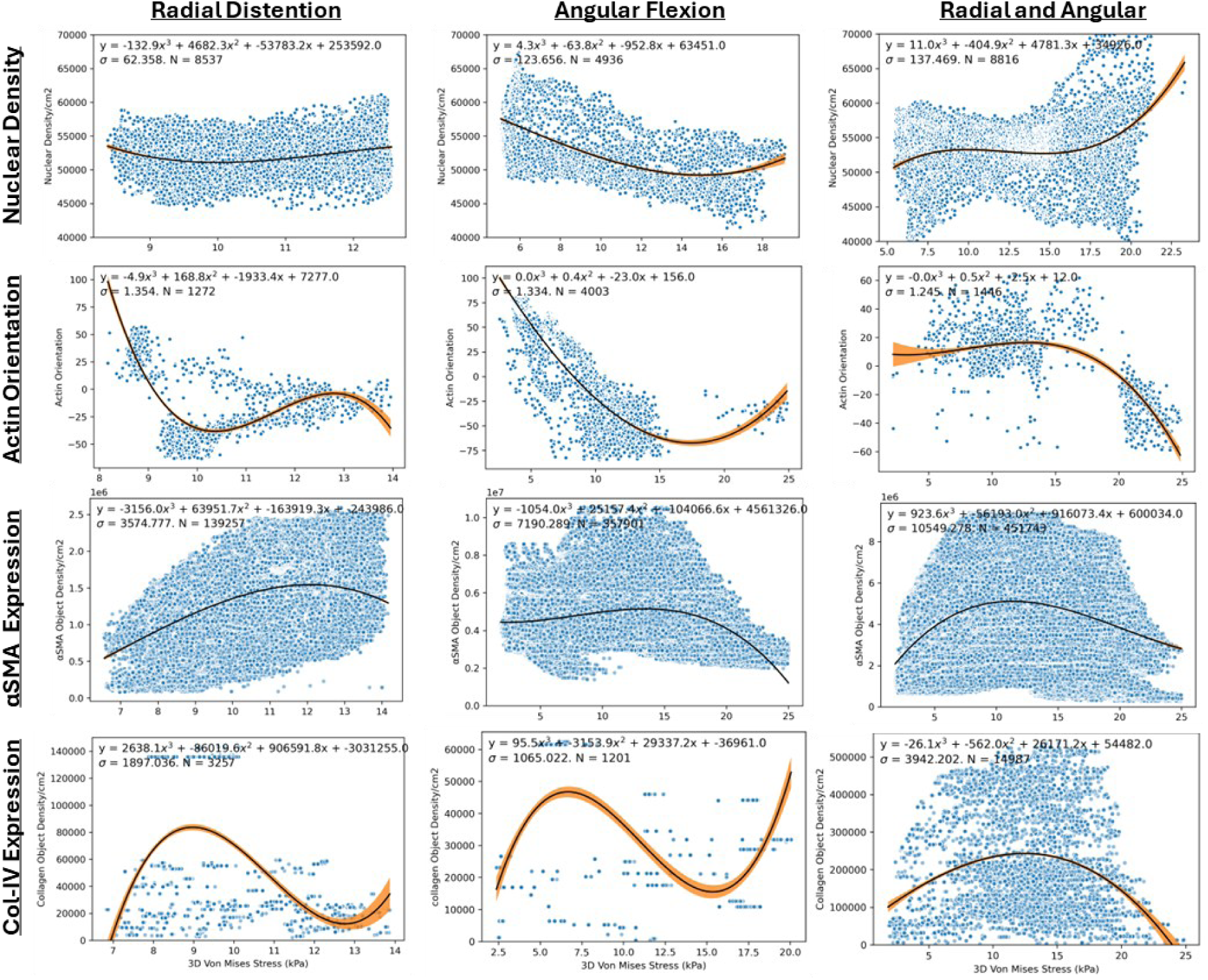
Impact of 3D Von Mises Stress on bigMACS Cell Behaviours. Scatterplots of 3D Von Mises Stress (kPa; x-axis) versus nuclear (DAPI stain) density, α-SMA stain density, and Col-IV stain density per cm^2^ or average actin orientation, as calculated in a 50 µm radius around each detected stain. Third-order polynomials were fitted to the imaging data with 95% confidence intervals calculated. Regression statistical replicates were calculated as the number of stains detected per image (1,201 to 451,749) and not imaging, experimental, or biological replicates.

Cell orientation, measured as cytoskeletal protein actin orientation, was randomly aligned for static soft robot bioreactors both on the microscale and across the full image (**Figure 3**). In contrast, mechanically- conditioned versus static bigMACS all expressed a larger number of brighter, longer, actin stains whose bulk orientation was increasingly aligned in a parallel direction to the major stress angle, especially for bioreactors with greater mechanical stress. Cell orientation for mechanically-conditioned bigMACS was consistent for the 𝑦-axis but varied along the 𝑥-axis, similar to how mechanical forces varied along 𝑥 but not 𝑦. Interestingly, cell orientation appeared to be correlated with moderate stress magnitude (**Figure 4**) and principal stress direction but only for radial bigMACS (**Figure 5**). Cell orientation within radial and angular conditioned bigMACS responded to changes of low-magnitude mechanical stress (5 – 10 kPa), though cell orientation on radial and angular combined bigMACS changed for stress magnitudes above 20 kPa (**Figure 4**). Cell orientation was inversely correlated to XY Mohr’s angle for radial BigMACS, indicating that cells aligned perpendicular to principal stress direction (**Figure 5**).

**Figure 5:**
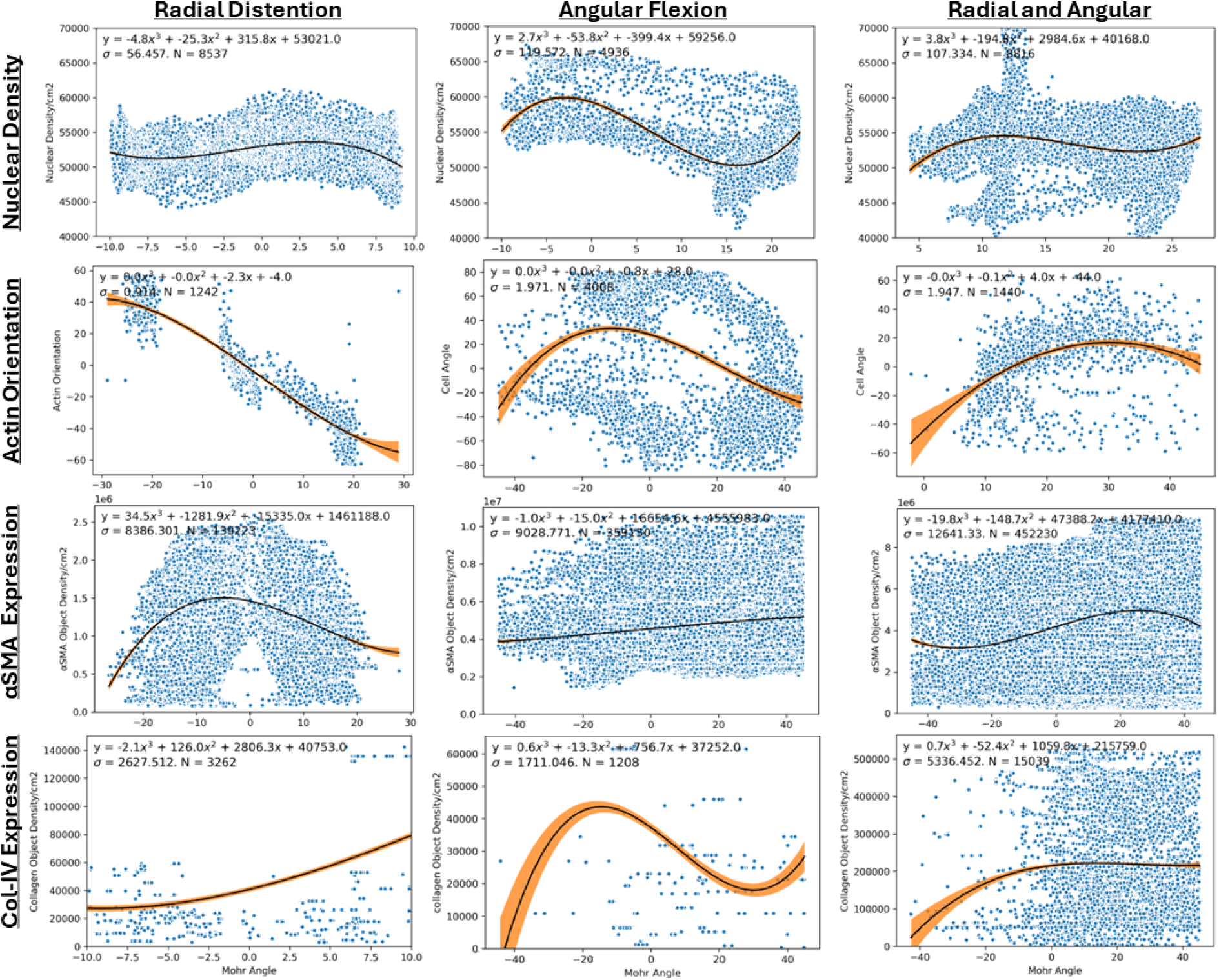
Impact of XY Mohr’s Stress Angle on bigMACS Cell Behaviours. Scatterplots of XY Mohr’s Stress Angle (x-axis) versus nuclear (DAPI stain) density, α-SMA stain density, and Col-IV stain density per cm^2^ or average actin orientation, as calculated in a 50 µm radius around each detected stain. Third-order polynomials were fitted to the imaging data with 95% confidence intervals calculated. Regression statistical replicates were calculated as the number of stains detected per image (1,201 to 451,749) and not imaging, experimental, or biological replicates.

Smooth muscle differentiation occurs as a consequence of MSC mechanical conditioning, even in the absence of media-supplemented soluble myogenic factors. Smooth muscle differentiation of MSCs is phenotypically detected by an upregulation in alpha smooth muscle actin (α-SMA). We identified all α-SMA stains in confocal images which fluoresced brighter than their respective isotype controls, and then calculated the local density of α-SMA around each α-SMA stain. While negligible α-SMA was detected in static bigMACS, 10-to-20 greater α-SMA was found in all three mechanically conditioned bigMACS (**Figure 3**). Mechanical stress magnitude was positively correlated with α-SMA stain density at lower 3D Von Mises Stress range (0 to 12 kPa) for all mechanically conditioned bigMACS, and then negatively correlated at higher 3D Von Mises Stresses (15 kPa to 25 kPa; **Figure 4**). This is in line with prior work suggesting super-physiological stress might shift cells to focus on survival or quiescence instead of smooth muscle differentiation [41, 42]. Expression of α-SMA in radial distension bigMACS was most intense near XY Mohr’s Stress Angles of 0°; however, this is likely because it is where mechanical stress magnitude was greatest in radial distension. Similar trends were not found between α-SMA and principal stress direction for other mechanical BigMACS conditions (**Figure 5**).

Matrix deposition of vascular collagen type 4 (Col-IV) from MSCs or SMCs, found in the basal lamina of arteries, is enhanced under mechanical stimulation [43]. Similar as α-SMA, we identified all Col-IV stains in confocal images which fluoresced brighter than their respective isotype controls, and then calculated the local density of Col-IV around each Col-IV stain. Col-IV was negligibly expressed in static bigMACS, more expressed in radial or angular conditioned bigMACS, and significantly more expressed in combined radial and angular bigMACS. No local Col-IV deposition trends were identified for radial or angular conditioning. For bigMACS with combined radial and angular conditioning, more Col-IV was deposited in the centre of the bigMACS, where there was an intermediate stress magnitude and positive principal stress direction (most bigMACS exhibited positive angles). While Col-IV was only deposited by MSCs in bigMACS conditioned with the highest stress magnitudes, this deposition seemed to occur across the whole culture chamber, and did not localise in regions of higher mechanical stress.

In summary, big mechanically active culture systems (bigMACS) are of increasing importance in tissue engineering and regenerative medicine. These bigMACS apply dynamic mechanical forces which reproduce our body’s natural fluid flow, locomotion, or flexion to model regeneration or disease, to distribute nutrients or growth factors for higher density cell growth, to trigger the continuous release of cell or cell-based products throughout culture (such as extracellular vesicles) [44–47], and, in this application, to promote stem cell proliferation or differentiation in the absence of exogenous factors.

Despite the increased use of bigMACS in disease modelling and in cell or tissue biomanufacturing, the effect of mechanical forces on cell behaviour is non-deterministic and poorly understood. Lessons from microscale mechanobiology and macroscale biomechanics have taught us that mechanical forces create genetic, phenotypic, and functional changes in single cells which exhibit paracrine signals that, together with mechanical forces, easily propagate across a confluent tissue [42, 48, 49]. It is challenging to study these two mechanisms using bigMACS as the fabrication of big culture platforms can often create artefacts at the single-cell microscale [50], such as 3D printed layer lines [51], and because the application of physiological mechanical forces vary over large-scale cell culture chambers.

## 4. Conclusion

We have previously engineered a bigMACS in the design of a soft robot bioreactor, able to mimic multi-axial forces present in the human femoropopliteal artery, and culture a confluent layer of mesenchymal stem cells (MSCs). We found increased mechanical conditioning forces led to increases in cell density, cell alignment, smooth muscle differentiation, and extracellular matrix deposition across the centimetre-scale culture reservoir. These outcomes have important consequences for disease modelling and cell or tissue biomanufacturing platforms.

In this study, we developed a computational modelling and quantitative histology framework to identify what local mechanical force is required for desired local and bulk changes in cell behaviour. If we could define these relationships, we could formulate the fundamental mechanical design equations for gradient tissue modelling and biomanufacturing. Due to microscale artefacts ubiquitous in bigMACS fabrication or surface processes, we developed our centimetre-scale FEA simulation to have microscale resolution. For each of the 2-to-10 thousand cells imaged in our bigMACS culture chamber, our FEA simulation would average hundreds of local mechanical stresses required for simulation convergence.

Our innovative FEA simulation and quantitative histology framework derived new relationships on how principal stress magnitude and direction appear to pattern MSC behaviour. Specifically, smooth muscle (SMC) differentiation by MSCs was found to have a positive correlation with local areas of high stress in all mechanically conditioned bigMACS, suggesting that soft robotic bioreactors are a promising tool to controllably design and engineer gradient smooth muscle tissue, including zones of potent MSCs and zones of differentiated SMCs. We found that cell orientation is inversely proportional to mechanical principal stress direction but only for radial BigMACS and not more complex actuation modes. In contrast, we found that increasing mechanical stress magnitude could not be used to locally pattern MSC nuclear density and Col-IV deposition; indicating these are cell behaviours that rapidly propagate across culture space and would be challenging to spatially confine with bigMACS. The local principal stress direction did not play any significant role on local cell orientation or other behaviours, suggesting some cell behaviours are dependent on regional mechanics and are challenging to locally control and pattern at a single-cell scale. The application of our framework to our soft robotic bigMACS bioreactor is limited by the analysis of only one culture replicate of each mechanical actuation condition. Biological and technical replicates must be preformed in future studies to validate any proposed mechanobiology design equations.

To our knowledge, this is the first FEA simulation able to predict subcellular mechanical forces in the presence of microroughness often found in dynamic tissue engineered cell cultures; and this is the first framework that applies these microscale FEA simulation results to single-cell quantitative histology across large, mechanically-patterned tissues (10,000’s of cells; centimetres of tissue). Our new FEA mechanical simulation and quantitative histology framework are a powerful tool for investigating how big mechanically active culture systems (bigMACS) can be designed or engineered to apply mechanical forces which control microscale gradients of mechanical forces able to locally or regionally pattern cells and tissues for disease modelling and biomanufacturing. This framework opens new doors to formulate the fundamental design equations of mechanically-patterned tissue biomanufacturing.

## Acknowledgements

This work was supported by a GOstralia!-GOzealand! GmbH Scholarship to SS, a Bionics Queensland Grand Challenge Award to SS, CAF, MCA, a British Heart Foundation Centre of Research Excellence Award (RE/18/4/34215) and a Delft Technology Fellowship to SP, a Strategic Partnership Award for Research Collaboration of the Chinese University of Hong Kong (4930640) to HFC and MCA, Australian Research Council funding to YCT and MCA (FT150100398, DP230100721, DE220100757), and an Advance Queensland fellowship (AQIRF1312018) to MCA.

## Supplementary Information

**Figure S1:**
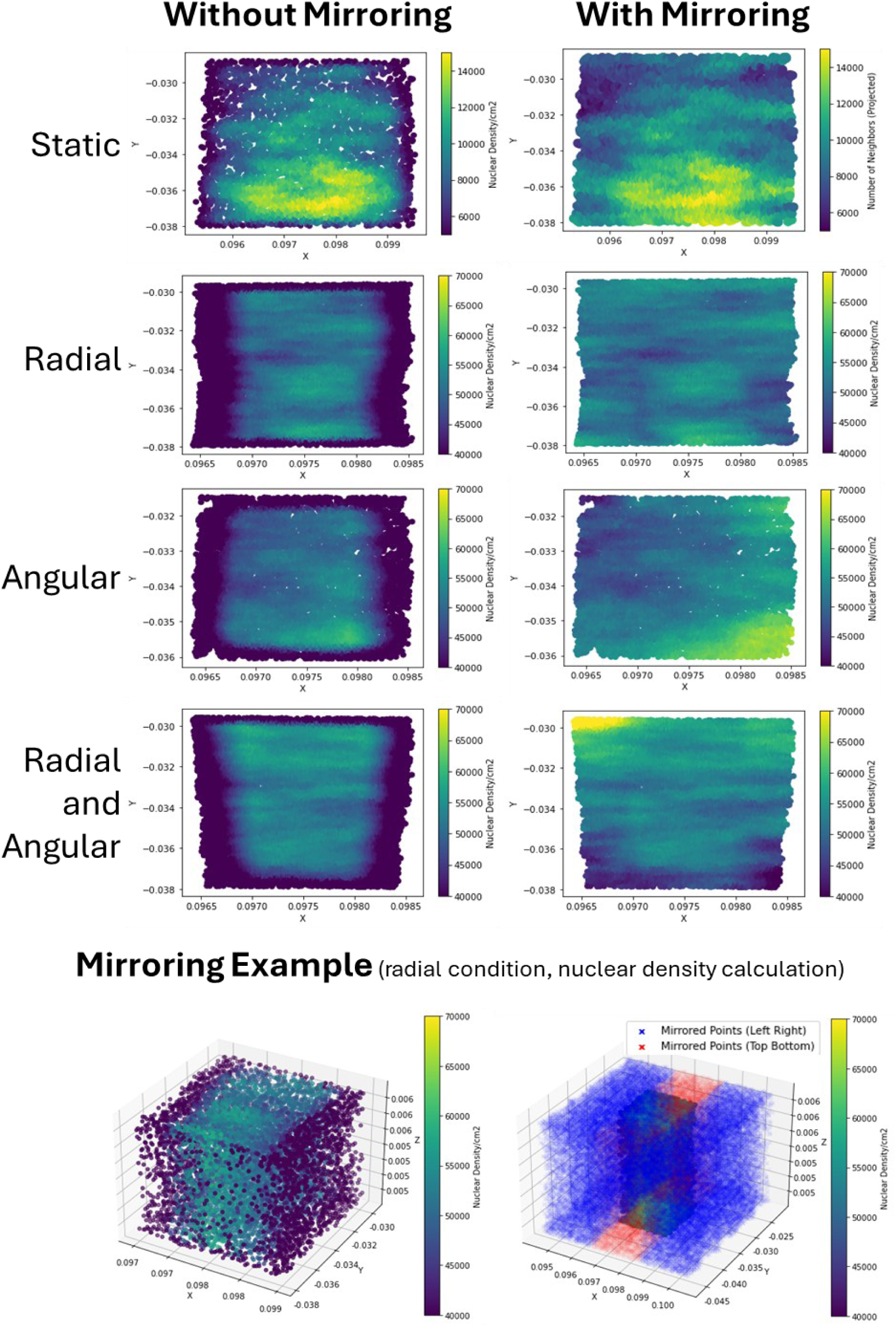
Original stain density calculations (left), and stain density calculations preformed after mirroring stain locations over the boundary of the tissue (right). Stain density calculation artefacts indicated on the ‘without mirroring’ column as dark edge regions were removed after mirroring.

**Figure S2:**
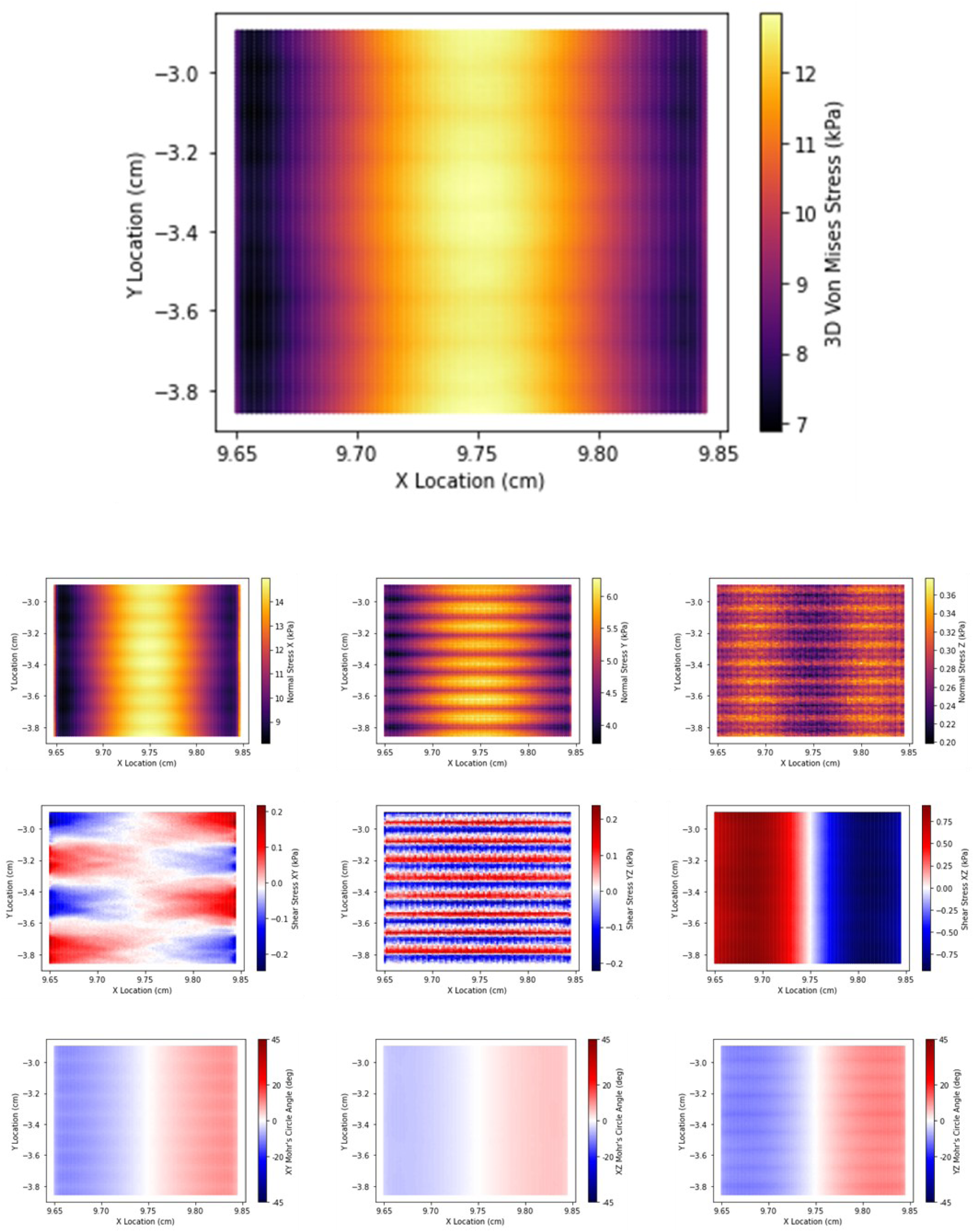
Summary of mechanical force outputs for radial bigMACS with microgrooves over its surface. The top graph represents the 3D Von Mises Stress. The second row of graphs represents three dimensions of 2D Von Mises Stress. The third row represents three dimensions of shear stresses. The fourth row represents three dimensions of 2D Mohr’s Angles.

**Figure S3:**
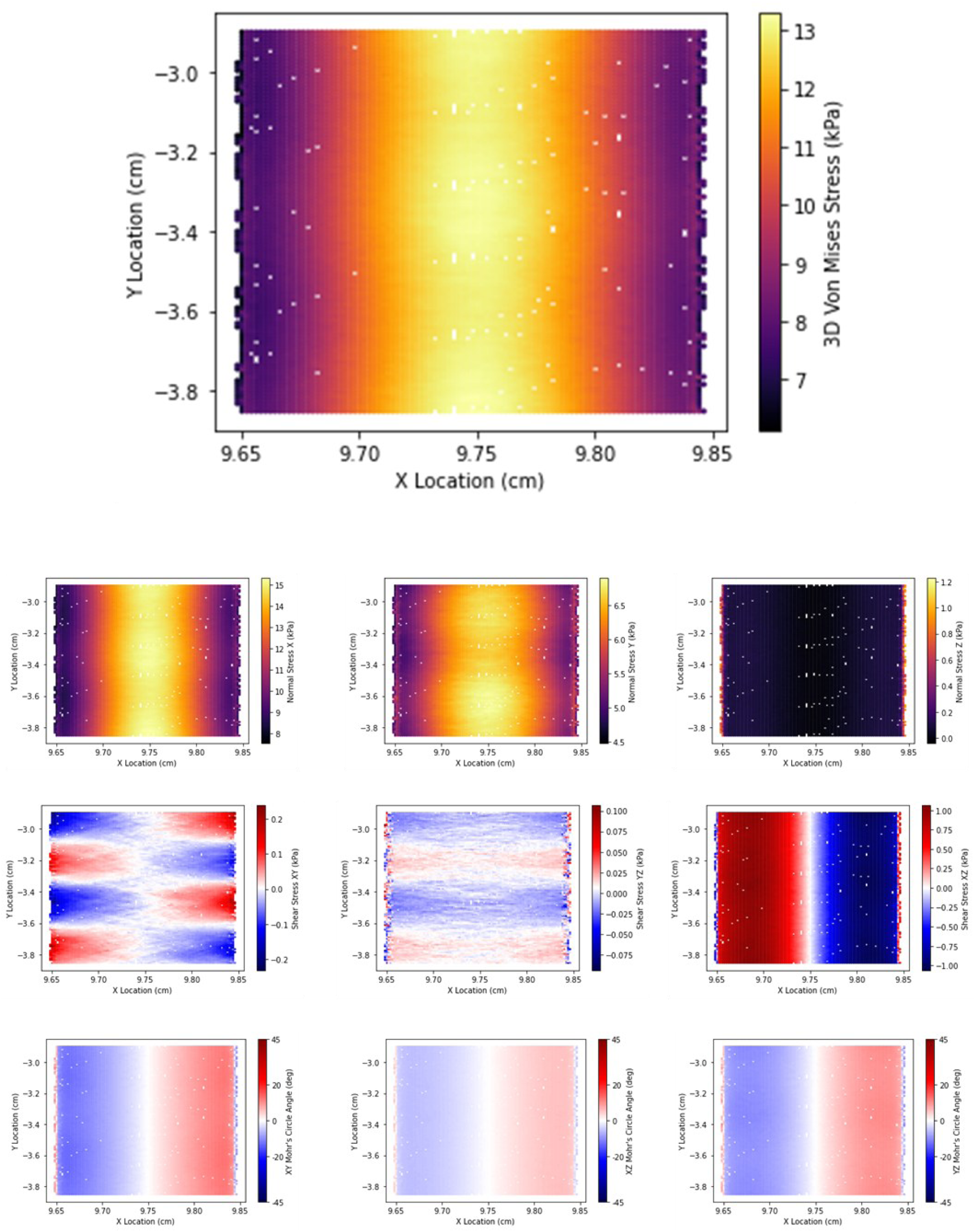
Summary of mechanical force outputs for radial bigMACS with a smooth surface. Smooth surface bigMACS were simulated with 100-fold fewer FEA nodes, a lower mesh density. The top graph represents the 3D Von Mises Stress. The second row of graphs represents three dimensions of 2D Von Mises Stress. The third row represents three dimensions of shear stresses. The fourth row represents three dimensions of 2D Mohr’s Angles.

**Figure S4:**
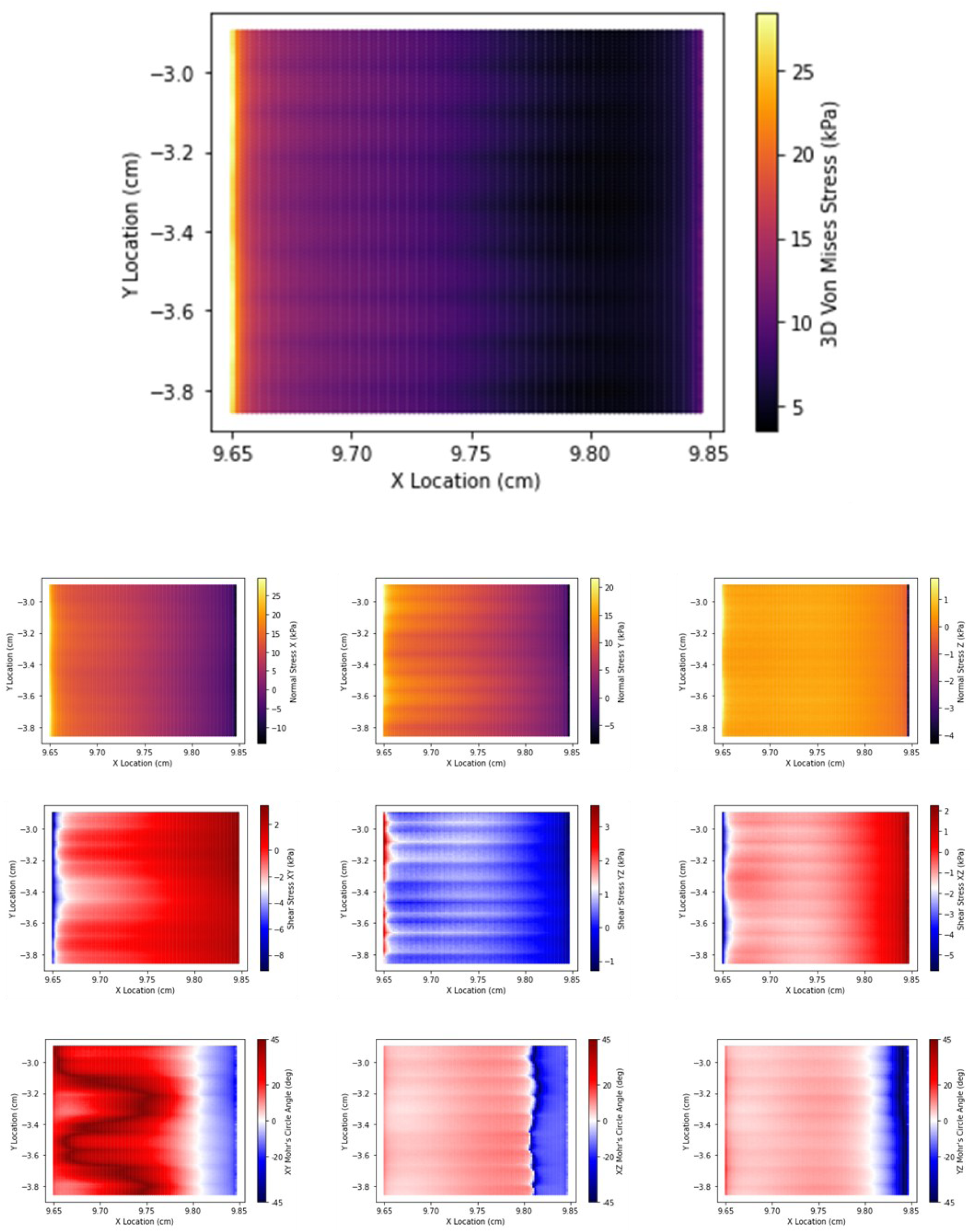
Summary of mechanical force outputs for angular bigMACS with microgrooves over its surface. The top graph represents the 3D Von Mises Stress. The second row of graphs represents three dimensions of 2D Von Mises Stress. The third row represents three dimensions of shear stresses. The fourth row represents three dimensions of 2D Mohr’s Angles.

**Figure S5:**
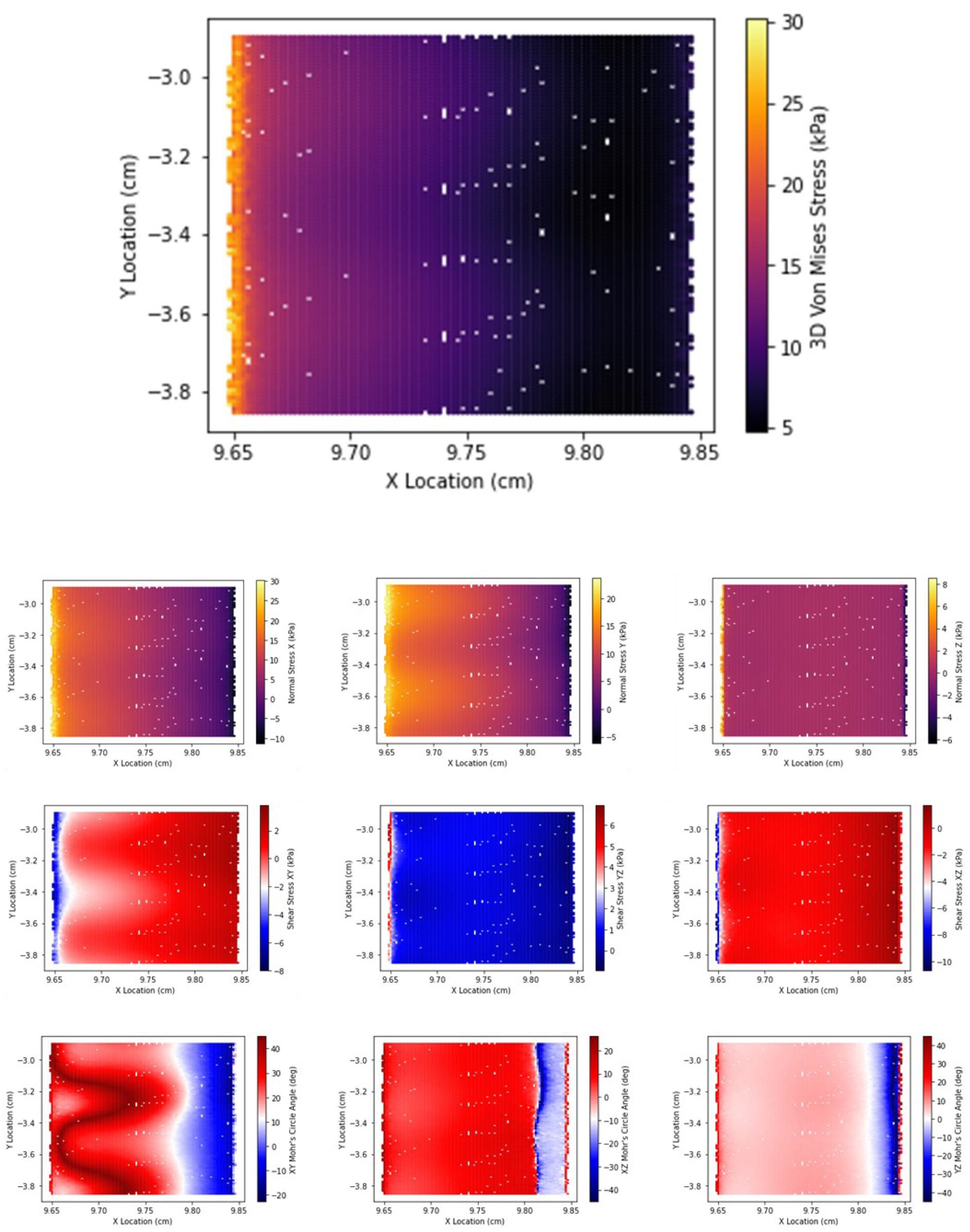
Summary of mechanical force outputs for angular bigMACS with a smooth surface. Smooth surface bigMACS were simulated with 100-fold fewer FEA nodes, a lower mesh density. The top graph represents the 3D Von Mises Stress. The second row of graphs represents three dimensions of 2D Von Mises Stress. The third row represents three dimensions of shear stresses. The fourth row represents three dimensions of 2D Mohr’s Angles.

**Figure S6:**
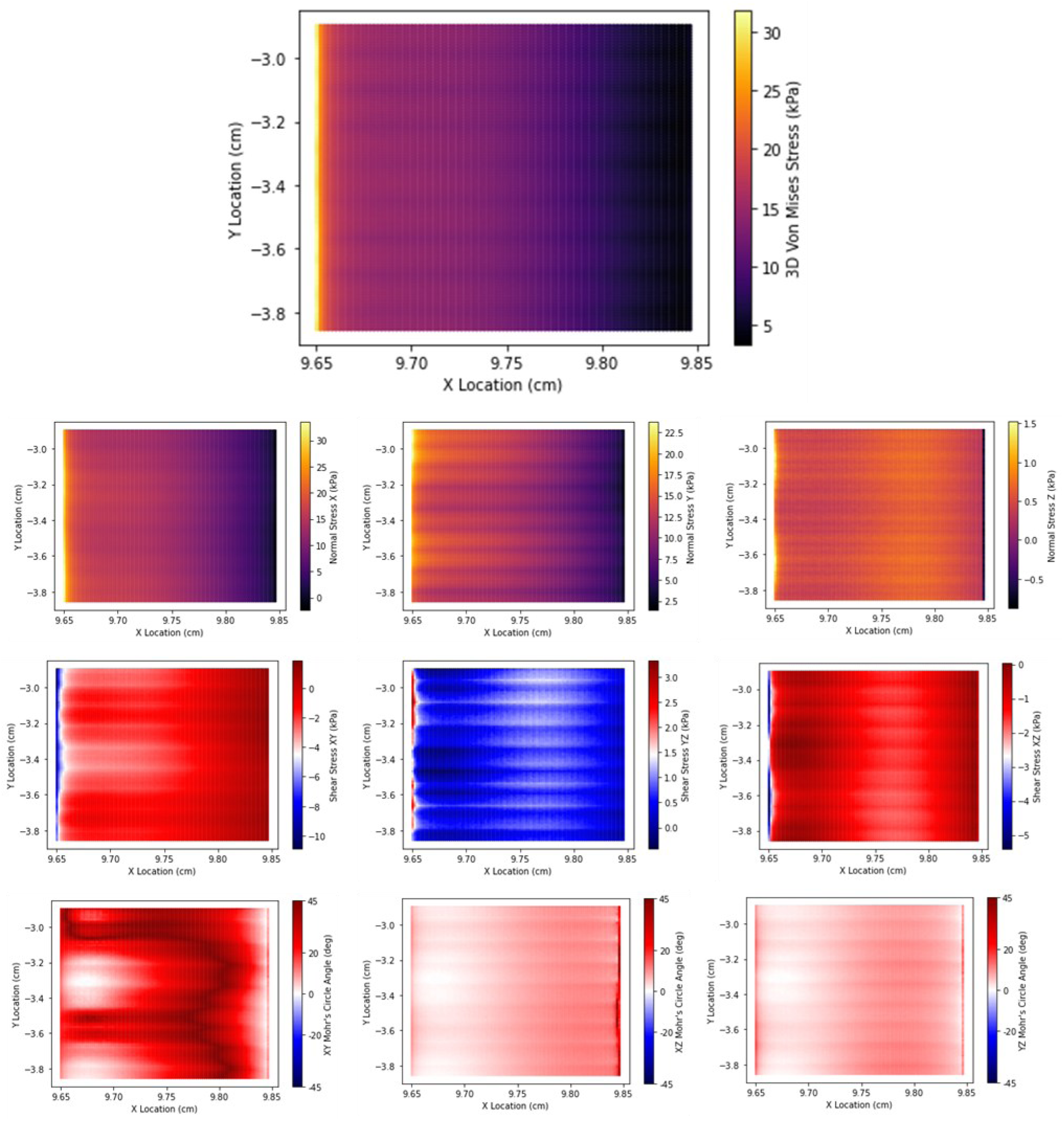
Summary of mechanical force outputs for combined (radial plus angular actuation) bigMACS with microgrooves over its surface. The top graph represents the 3D Von Mises Stress. The second row of graphs represents three dimensions of 2D Von Mises Stress. The third row represents three dimensions of shear stresses. The fourth row represents three dimensions of 2D Mohr’s Angles.

**Figure S7:**
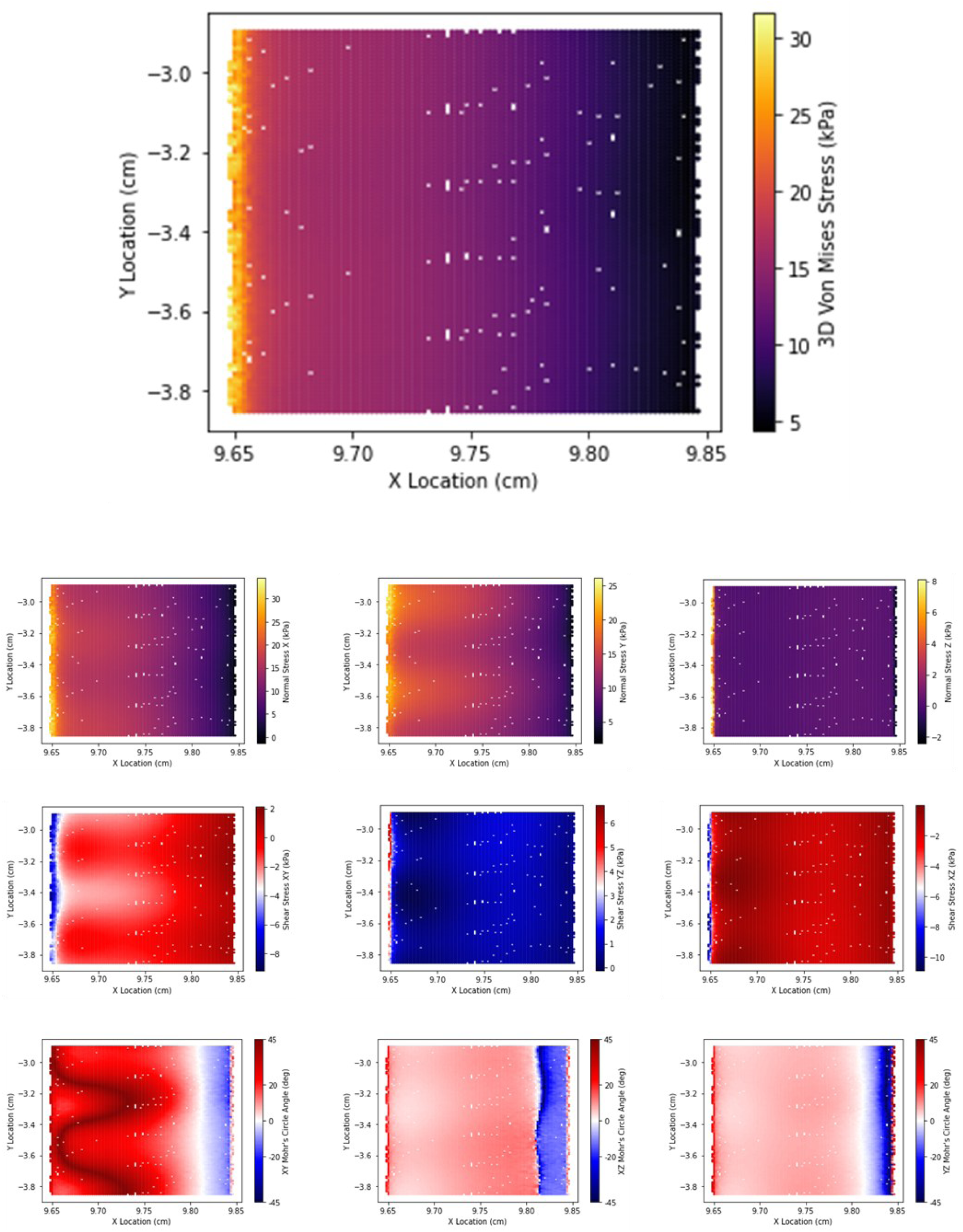
Summary of mechanical force outputs for combined (radial and angular actuation) bigMACS with a smooth surface. Smooth surface bigMACS were simulated with 100-fold fewer FEA nodes, a lower mesh density. The top graph represents the 3D Von Mises Stress. The second row of graphs represents three dimensions of 2D Von Mises Stress. The third row represents three dimensions of shear stresses. The fourth row represents three dimensions of 2D Mohr’s Angles.

## References

[1] K. H. Vining and D. J. Mooney, "Mechanical forces direct stem cell behaviour in development and regeneration," Nat Rev Mol Cell Biol, vol. 18, pp. 728–742, Dec 2017.

[2] M. Sun, G. Chi, P. Li, S. Lv, J. Xu, Z. Xu, et al., "Effects of Matrix Stiffness on the Morphology, Adhesion, Proliferation and Osteogenic Differentiation of Mesenchymal Stem Cells," Int J Med Sci, vol. 15, pp. 257–268, 2018.

[3] S. Kim, M. Uroz, J. L. Bays, and C. S. Chen, "Harnessing Mechanobiology for Tissue Engineering," Dev Cell, vol. 56, pp. 180–191, Jan 25 2021.

[4] K. Katoh, "Effects of Mechanical Stress on Endothelial Cells In Situ and In Vitro," Int J Mol Sci, vol. 24, Nov 20 2023.

[5] X. Zhang, S. Zhang, and T. Wang, "How the mechanical microenvironment of stem cell growth affects their differentiation: a review," Stem Cell Research & Therapy, vol. 13, p. 415, 2022/08/13 2022.

[6] X. Huang, Z. Huang, W. Gao, W. Gao, R. He, Y. Li, et al., "Current Advances in 3D Dynamic Cell Culture Systems," Gels, vol. 8, p. 829, 2022.

[7] K. Uto, J. H. Tsui, C. A. DeForest, and D. H. Kim, "Dynamically Tunable Cell Culture Platforms for Tissue Engineering and Mechanobiology," Prog Polym Sci, vol. 65, pp. 53–82, Feb 2017.

[8] D. A. Gaspar, V. Gomide, and F. J. Monteiro, "The role of perfusion bioreactors in bone tissue engineering," Biomatter, vol. 2, pp. 167-75, Oct-Dec 2012.

[9] S. M. Somers, A. A. Spector, D. J. DiGirolamo, and W. L. Grayson, "Biophysical Stimulation for Engineering Functional Skeletal Muscle," Tissue Eng Part B Rev, vol. 23, pp. 362-372, Aug 2017.

[10] J. Paek, J. W. Song, E. Ban, Y. Morimitsu, C. O. Osuji, V. B. Shenoy, and D. D. Huh, "Soft robotic constrictor for in vitro modeling of dynamic tissue compression," Scientific Reports, vol. 11, p. 16478, 2021/08/13 2021.

[11] Y. Sun, B. Wan, R. Wang, B. Zhang, P. Luo, D. Wang, et al., "Mechanical stimulation on mesenchymal stem cells and surrounding microenvironments in bone regeneration: regulations and applications," Frontiers in cell and developmental biology, vol. 10, p. 808303, 2022.

[12] A. J. Steward and D. J. Kelly, "Mechanical regulation of mesenchymal stem cell differentiation," Journal of Anatomy, vol. 227, pp. 717–731, 2015.

[13] Y.-J. Chen, C.-H. Huang, I. C. Lee, Y.-T. Lee, M.-H. Chen, and T.-H. Young, "Effects of Cyclic Mechanical Stretching on the mRNA Expression of Tendon/Ligament-Related and Osteoblast-Specific Genes in Human Mesenchymal Stem Cells," Connective Tissue Research, vol. 49, pp. 7–14, 2008/01/01 2008.

[14] R. Torii, R. I. Velliou, D. Hodgson, and V. Mudera, "Modelling multi-scale cell-tissue interaction of tissue-engineered muscle constructs," J Tissue Eng, vol. 9, p. 2041731418787141, Jan-Dec 2018.

[15] C. S. Chen, J. Tan, and J. Tien, "Mechanotransduction at Cell-Matrix and Cell-Cell Contacts," Annual Review of Biomedical Engineering, vol. 6, pp. 275–302, 2004.

[16] Y. Hou, L. Yu, W. Xie, L. C. Camacho, M. Zhang, Z. Chu, et al., "Surface Roughness and Substrate Stiffness Synergize To Drive Cellular Mechanoresponse," Nano Letters, vol. 20, pp. 748–757, 2020/01/08 2020.

[17] S. Mok, S. Al Habyan, C. Ledoux, W. Lee, K. N. MacDonald, L. McCaffrey, and C. Moraes, "Mapping cellular-scale internal mechanics in 3D tissues with thermally responsive hydrogel probes," Nature Communications, vol. 11, p. 4757, 2020/09/21 2020.

[18] J. S. Park, J. S. F. Chu, C. Cheng, F. Chen, D. Chen, and S. Li, "Differential effects of equiaxial and uniaxial strain on mesenchymal stem cells," Biotechnology and Bioengineering, vol. 88, pp. 359–368, 2004.

[19] M.-M. Khani, M. Tafazzoli-Shadpour, Z. Goli-Malekabadi, and N. Haghighipour, "Mechanical characterization of human mesenchymal stem cells subjected to cyclic uniaxial strain and TGF-β1," Journal of the Mechanical Behavior of Biomedical Materials, vol. 43, pp. 18–25, 2015/03/01/ 2015.

[20] M. Koike, H. Shimokawa, Z. Kanno, K. Ohya, and K. Soma, "Effects of mechanical strain on proliferation and differentiation of bone marrow stromal cell line ST2," Journal of Bone and Mineral Metabolism, vol. 23, pp. 219–225, 2005/05/01 2005.

[21] M. Jagodzinski, M. Drescher, J. Zeichen, S. Hankemeier, C. Krettek, U. Bosch, and M. Van Griensven, "Effects of cyclic longitudinal mechanical strain and dexamethasone on osteogenic differentiation of human bone marrow stromal cells," Eur Cell Mater, vol. 7, p. 41, 2004.

[22] R. D. Sumanasinghe, S. H. Bernacki, and E. G. Loboa, "Osteogenic Differentiation of Human Mesenchymal Stem Cells in Collagen Matrices: Effect of Uniaxial Cyclic Tensile Strain on Bone Morphogenetic Protein (BMP-2) mRNA Expression," Tissue Engineering, vol. 12, pp. 3459–3465, 2006.

[23] A. Charoenpanich, M. E. Wall, C. J. Tucker, D. M. K. Andrews, D. S. Lalush, and E. G. Loboa, "Microarray Analysis of Human Adipose-Derived Stem Cells in Three-Dimensional Collagen Culture: Osteogenesis Inhibits Bone Morphogenic Protein and Wnt Signaling Pathways, and Cyclic Tensile Strain Causes Upregulation of Proinflammatory Cytokine Regulators and Angiogenic Factors," Tissue Engineering Part A, vol. 17, pp. 2615-2627, 2011.

[24] Y. Qiu, J. Lei, T. J. Koob, and J. S. Temenoff, "Cyclic tension promotes fibroblastic differentiation of human MSCs cultured on collagen-fibre scaffolds," Journal of Tissue Engineering and Regenerative Medicine, vol. 10, pp. 989–999, 2016.

[25] P. J. Yang, M. E. Levenston, and J. S. Temenoff, "Modulation of Mesenchymal Stem Cell Shape in Enzyme-Sensitive Hydrogels Is Decoupled from Upregulation of Fibroblast Markers Under Cyclic Tension," Tissue Engineering Part A, vol. 18, pp. 2365–2375, 2012/11/01 2012.

[26] C. Fell, T. L. Brooks-Richards, M. A. Woodruff, and M. C. Allenby, "Soft pneumatic actuators for mimicking multi-axial femoropopliteal artery mechanobiology," Biofabrication, vol. 14, p. 035005, 2022/04/20 2022.

[27] L. Lévesque, C. Loy, A. Lainé, B. Drouin, P. Chevallier, and D. Mantovani, "Incrementing the frequency of dynamic strain on SMC-cellularised collagen-based scaffolds affects extracellular matrix remodeling and mechanical properties," ACS Biomaterials Science & Engineering, vol. 4, pp. 3759–3767, 2017.

[28] M. Rabbani, M. Tafazzoli-Shadpour, M. A. Shokrgozar, M. Janmaleki, and M. Teymoori, "Cyclic stretch effects on adipose-derived stem cell stiffness, morphology and smooth muscle cell gene expression," Tissue Engineering and Regenerative Medicine, vol. 14, pp. 279–286, 2017.

[29] B. Walters, T. Uynuk-Ool, M. Rothdiener, J. Palm, M. L. Hart, J. P. Stegemann, and B. Rolauffs, "Engineering the geometrical shape of mesenchymal stromal cells through defined cyclic stretch regimens," Scientific reports, vol. 7, p. 6640, 2017.

[30] J. Kreutzer, M. Viehrig, R.-P. Pölönen, F. Zhao, M. Ojala, K. Aalto-Setälä, and P. Kallio, "Pneumatic unidirectional cell stretching device for mechanobiological studies of cardiomyocytes," Biomechanics and Modeling in Mechanobiology, vol. 19, pp. 291–303, 2020/02/01 2020.

[31] P. Han, T. Guo, A. Jayasree, G. A. Gomez, K. Gulati, and S. Ivanovski, "Tunable nano-engineered anisotropic surface for enhanced mechanotransduction and soft-tissue integration," Nano Research, vol. 16, pp. 7293–7303, 2023/05/01 2023.

[32] R. M. Hackett, Hyperelasticity primer: Springer, 2016.

[33] R. W. Ogden, "Large deformation isotropic elasticity–on the correlation of theory and experiment for incompressible rubberlike solids," Proceedings of the Royal Society of London. A. Mathematical and Physical Sciences, vol. 326, pp. 565–584, 1972.

[34] P. Moseley, J. M. Florez, H. A. Sonar, G. Agarwal, W. Curtin, and J. Paik, "Modeling, design, and development of soft pneumatic actuators with finite element method," Advanced engineering materials, vol. 18, pp. 978–988, 2016.

[35] [35] The University of Queensland Research Computing Centre. (2024). Bunya supercomputer. Available: 10.48610/wf6c-qy55

[36] P. D. Barsanescu and A. M. Comanici, "von Mises hypothesis revised," Acta Mechanica, vol. 228, pp. 433–446, 2017/02/01 2017.

[37] J. Y. Lee, H. R. Ryu, and Y. T. Park, "Finite element implementation for computer-aided education of structural mechanics: Mohr’s circle and its practical use," Computer Applications in Engineering Education, vol. 22, pp. 494–508, 2014.

[38] O. Mohr, "Über die Darstellung des Spannungszustandes und des Deformationszustandes eines Körperelementes und über die Anwendung derselben in der Festigkeit," Civilingenieur, vol. 28, 1854.

[39] R. Rezakhaniha, A. Agianniotis, J. T. Schrauwen, A. Griffa, D. Sage, C. V. Bouten, et al., "Experimental investigation of collagen waviness and orientation in the arterial adventitia using confocal laser scanning microscopy," Biomech Model Mechanobiol, vol. 11, pp. 461-73, Mar 2012.

[40] M. C. Allenby, R. Misener, N. Panoskaltsis, and A. Mantalaris, "A quantitative three-dimensional image analysis tool for maximal acquisition of spatial heterogeneity data," Tissue Engineering Part C: Methods, vol. 23, pp. 108–117, 2017.

[41] F. Relaix, M. Bencze, M. J. Borok, A. Der Vartanian, F. Gattazzo, D. Mademtzoglou, et al., "Perspectives on skeletal muscle stem cells," Nature Communications, vol. 12, p. 692, 2021/01/29 2021.

[42] J. Tao, M. I. Choudhury, D. Maity, T. Kim, S. X. Sun, and C.-M. Fan, "Mechanical compression creates a quiescent muscle stem cell niche," Communications Biology, vol. 6, p. 43, 2023/01/13 2023.

[43] T. Sillat, R. Saat, R. Pöllänen, M. Hukkanen, M. Takagi, and Y. T. Konttinen, "Basement membrane collagen type IV expression by human mesenchymal stem cells during adipogenic differentiation," J Cell Mol Med, vol. 16, pp. 1485-95, Jul 2012.

[44] W. Cao, W. Lin, H. Cai, Y. Chen, Y. Man, J. Liang, et al., "Dynamic mechanical loading facilitated chondrogenic differentiation of rabbit BMSCs in collagen scaffolds," Regenerative Biomaterials, vol. 6, pp. 99–106, 2019.

[45] F. Wei, K. Flowerdew, M. Kinzel, L. E. Perotti, J. Asiatico, M. Omer, et al., "Changes in interstitial fluid flow, mass transport and the bone cell response in microgravity and normogravity," Bone Research, vol. 10, p. 65, 2022/11/21 2022.

[46] V. Tsata and D. Beis, "In Full Force. Mechanotransduction and Morphogenesis during Homeostasis and Tissue Regeneration," J Cardiovasc Dev Dis, vol. 7, Oct 1 2020.

[47] T. Y. Wong, S. N. Chang, R. C. Jhong, C. J. Tseng, G. C. Sun, and P. W. Cheng, "Closer to Nature Through Dynamic Culture Systems," Cells, vol. 8, Aug 21 2019.

[48] C. Argentati, F. Morena, I. Tortorella, M. Bazzucchi, S. Porcellati, C. Emiliani, and S. Martino, "Insight into Mechanobiology: How Stem Cells Feel Mechanical Forces and Orchestrate Biological Functions," Int J Mol Sci, vol. 20, Oct 26 2019.

[49] Y. Sun, B. Wan, R. Wang, B. Zhang, P. Luo, D. Wang, et al., "Mechanical Stimulation on Mesenchymal Stem Cells and Surrounding Microenvironments in Bone Regeneration: Regulations and Applications," Frontiers in Cell and Developmental Biology, vol. 10, 2022-January-21 2022.

[50] A. Madrid-Sánchez, F. Duerr, Y. Nie, H. Thienpont, and H. Ottevaere, "Fabrication of large-scale scaffolds with microscale features using light sheet stereolithography," Int J Bioprint, vol. 9, p. 650, 2023.

[51] M. J. Lerman, J. Lembong, G. Gillen, and J. P. Fisher, "3D printing in cell culture systems and medical applications," Appl Phys Rev, vol. 5, p. 041109, Dec 2018.

